# Space, time, and captivity: quantifying the factors influencing the fecal microbiome of an alpine ungulate

**DOI:** 10.1101/475459

**Authors:** Sarah E. Haworth, Kevin S. White, Steeve D. Côté, Aaron B.A. Shafer

## Abstract

The community of microorganisms in the gut is affected by host species, diet, and environment and is linked to normal functioning of the host organism. Although the microbiome fluctuates in response to host demands and environmental changes, there are core groups of microorganisms that remain relatively constant throughout the hosts lifetime. Ruminants are mammals that rely on highly specialized digestive and metabolic modifications, including microbiome adaptations, to persist in extreme environments. Here, we assayed the fecal microbiome of four mountain goat (*Oreamnos americanus*) populations in western North America. We quantified fecal microbiome diversity and composition among groups in the wild and captivity, across populations, and in a single group over time. There were no differences in community evenness or diversity across groups, although we observed a decreasing diversity trend across summer months. Pairwise sample estimates grouped the captive population distinctly from the wild populations, and moderately grouped the southern wild group distinctly from the two northern wild populations. We identified 33 genera modified by captivity, with major differences in key groups associated with cellulose degradation that likely reflect differences in diet. Our findings are consistent with other ruminant studies and provide baseline microbiome data in this enigmatic species, offering valuable insights into the health of wild alpine ungulates.

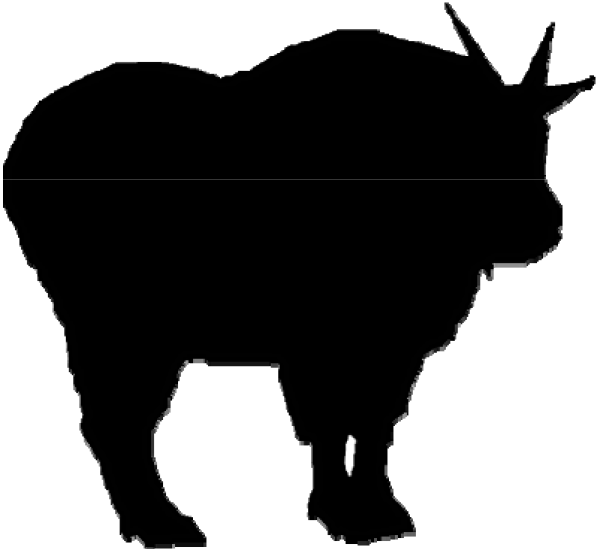

**Summary:** This study characterizes the microbiome of mountain goat (*Oreamnos americanus*) populations across populations and over summer months; we also quantified the effects of captivity to offer more insights into the health of alpine wildlife.

## Introduction

The microbiome describes a dynamic community of microorganisms that colonize organisms from birth onwards. Microorganisms in the gut play a key role in host physiological and immunological development (Berg, 1996), with fecal matter containing microbiome DNA that is shed during the digestion and egestion processes (Ingala et al., 2018; Vandeputte et al., 2016). The microbiome can vary according to the host species, age, diet, health, reproductive status, and external environment, but is directly linked to host health, including metabolism, immunity, and development (Nishida & Ochman, 2017). The fecal microbiome is reflective of a transient response to changes in the host, such as responses to dietary shifts across seasons, but core groups of microorganisms are found in stable relative abundances throughout the life of the host and the relative proportions of these groups can act like a signature of the host’s health and environment, with reductions in diversity associated closely with reductions in fitness (Amato et al., 2018; Clayton et al., 2018; Donaldson et al., 2015). The stable relative abundances of the core groups are directly related to the function and the demand of the core groups. The relative ratio of *Firmicutes* to *Bacteroidetes*, for one example, the two most commonly dominant core groups in mammal fecal microbiomes, can be used to discern between carnivorous and herbivorous mammals since each group is responsible for different metabolic demands (Huntington et al., 2019; Kreisinger et al., 2018; Muegge et al., 2011).

Measuring signatures in the fecal microbiome over time, between populations, such as between captive and wild populations of mammals, can help effectively monitor animal health (Bahrndorff et al., 2016; Bik et al., 2016). For example, many captive populations of mammal species suffer from poor health and some conditions have been correlated to fecal microbiome dysbiosis and decreased diversity microbial groups responsible for normal health (Clayton et al., 2018; Li et al., 2018; McKenzie et al., 2017). Actions that target the correction of the microbiome, meaning promoting signatures of core groups that are closer to what is seen in relevant wild and healthy populations, have been suggested as a feasible approach to help alleviate and manage health issues (Li et al., 2018). Moreover, evidence is mounting that supports the link between host phenotypic and genomic variation, and the microbiome (Carthey et al., 2018; Sharpton, 2018); this interaction is particularly relevant when it comes to rapidly changing environments. For example, the host microbial community of coral was critical in facilitating adaptation to warming temperatures (Webster et al., 2017; Ziegler et al. 2017).

Ruminants are herbivorous hooved mammals (ungulates) with specialized anatomical and physiological adaptations to accommodate the cellulolytic fermentation of low-nutrition, high-fiber plant materials (Noel et al., 2017; de Tarso et al., 2016). Numerous extant ruminant populations have been domesticated (e.g. cow, goat), while many of the remaining wild ruminant species are facing population declines that are directly and indirectly driven by human activity, including agricultural-related activities that have promoted predation, disease and parasitism expansions, environmental change, and have led to significant habitat loss (Smith et al., 2016; Di Marco et al., 2014; Martin, 2001). Non-invasive monitoring of the fecal microbiome of ruminant populations serves as a potential tool that can be used to inform management and conservation decisions aimed at improving the health of captive animals, promoting adaptation to environmental change, and preventing disease and parasitic outbreaks (McKenzie et al. 2017; Amato et al., 2013).

The North American mountain goat (*Oreamnos americanus*) is a sub-alpine ruminant and a symbolic icon of mountain wilderness. Mountain goats are found in fragmented, and occasionally isolated, sub-populations across the Western Cordillera mountain ranges (Shafer et al., 2012; Festa-Bianchet, 2008). Mountain goats show mixed seasonal and sexual habitat selection preferences with relatively small home ranges and limited inter- and intra-population movement, which possibly lends to low genetic variability (Shafer et al., 2012; Ortego et al., 2011; Poole & Heard, 2003). Their longer lifespans, upwards of 12 years, along with seasonally and sexually dependent habitat selection, makes mountain goats a unique study system for attempting to understand how variation in the microbiome supports their highly adapted and unique alpine lifestyle. Furthermore, the longevity of mountain goats, relative to many model and non-model systems, creates opportunities for researchers to conduct prolonged surveillance and infer individual trends over time (e.g. season, year), including non-invasive fecal microbiome studies.

Here we assayed the fecal microbiome of four western North American mountain goat populations, of which one was a captive population (Figure 1). We tested two hypotheses: i) captivity reduces microbial diversity in mammals (McKenzie et al., 2017; Sun et al., 2016; Kohl et al., 2014; Carey et al., 2013); and ii) diversity is negatively correlated to latitude (Dikongué et al., 2017). We further took advantage of temporal sampling in one population to assay changes over three summer months, where we hypothesized a shift in fecal microbiome diversity reflecting the decreased abundance of food resources over the summertime as resources are consumed and depleted (Hicks et al., 2018; Hu et al., 2018; Noel et al., 2017; Festa-Bianchet & Côté, 2008; Hamel & Côté, 2007). Collectively, we predicted that the effect of captivity would drive the strongest differences in the diversity and the composition in mountain goat fecal microbiomes, followed by seasonality and biogeography. This is the first study to establish a fecal microbiome profile for both captive and wild mountain goat populations and contributes valuable baseline knowledge on the diversity of microbiomes for different species (*sensu* McKenzie et al. 2017), while quantifying the influence of three important drivers of diversity and composition: space, time and, captivity.

**Figure 1.**
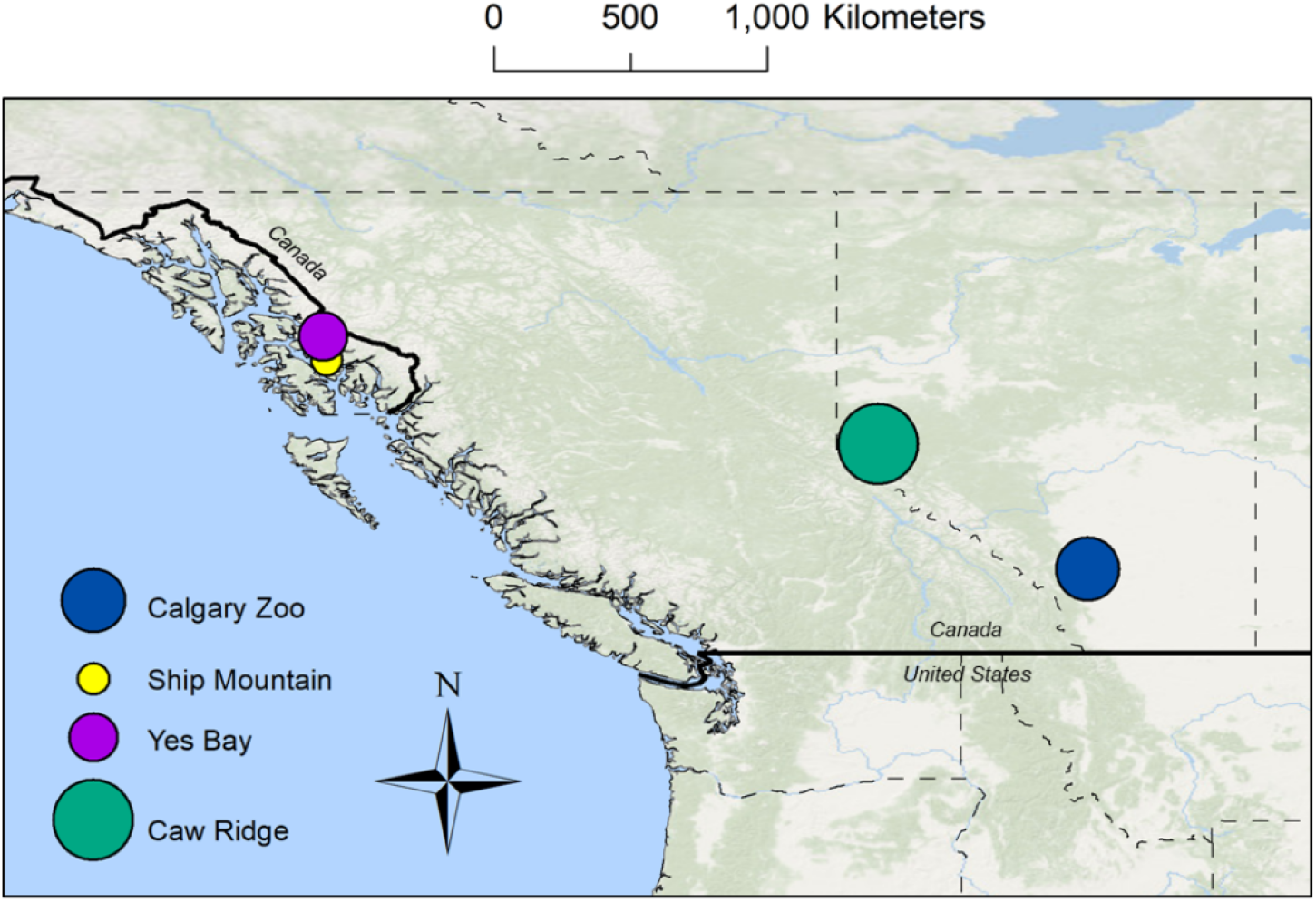
Geographical distribution of study sites in Alaska and Canada. The size of the points reflects the relative number of samples that originate from the study site.

## Methods

### Study species

Mountain goats are part of the Bovidae family and inhabit the mountains of western North America. Mountain goats are an important game species and populations are managed for harvest and non-consumptive uses. Mountain goats can be described as generalist foragers relative to other ruminant species, but it has been shown that they do show preferential selection of forage under some circumstances (Côté and Festa-Bianchet, 2003). During their daily movements, mountain goats spread an abundant amount of compact, dry pellets (feces) multiple times per day; such pellets are well preserved in the alpine environment they occupy.

### Study sites

The Calgary Zoo, Alberta, Canada (CZ; 51°N, 114°W) is one of Canada’s oldest accredited zoos. The zoo houses a small group of captive mountain goats that are fed year-round an *ad libidum* diet of herbivore pellets and mixed hay (see Table S1 for detailed diet information). This group of mountain goats is accessible for research and veterinary purposes. Caw Ridge, Alberta (CR; 54N, 119W) is an alpine and sub-alpine habitat in western Canada with a regional population of wild mountain goats that have been extensively monitored since 1989 (Ortego et al., 2011; Mainguy, Côté, & Coltman, 2009; Festa-Bianchet & Côté, 2008). As of 2017, CR was home to about 30 marked mountain goat individuals. This landscape only provides high quantities of quality forage during the summer months (June-August), while the rest of the year it is largely covered in snow or ice and forage availability is limited (Festa-Bianchet & Cote, 2008). Altitudinal migration does not occur in this population. Ship Mountain and Yes Bay (YB; 56N, −132W) each harbor relatively small (n = 40-50 individuals), geographically isolated populations located along the Cleveland Peninsula, southeastern Alaska, United States. Mountain goats on the Cleveland Peninsula are native and are genetically and morphologically distinct from surrounding mainland populations (Breen et al., submitted). Notably, mountain goats from this region are influenced by a maritime climate and seasonally migrate to low elevations during winter (Fox & Smith, 1988; Smith & Raedeke, 1982). Available diet information for the sampled populations is found in Supplementary Table S1.

### Sample collection and DNA extraction

Fresh mountain goat fecal samples (~10 pellets) were collected from CZ (August 2016), SM (July 2016), and YB (July 2016). Larger samples (>10 pellets) were collected from CR during the summer months (June-August 2016). Samples were collected at CR after observing groups of mountain goats defecate, whereas those in Alaska were collected by searching areas mountain goats were observed that day. Mountain goats typically defecate in defined, non-overlapping pellet groups and we only selected samples that were clearly from a single individual. Due to the nature of the sampling we were unable to sex or age the animal that defecated. The cool, alpine climate of the sampling environment naturally preserves samples and thus we do not suspect environmental contamination was a major factor. All samples (N=54), except those from CR, were stored immediately in 70% ethanol (EtOH) and at −20°C. CR samples were stored individually in plastic sample bags at −20°C. All surfaces were sterilized with 90% EtOH and 10% bleach solution to prevent environmental contamination. A small portion of a single fecal sample (~1/4 including exterior and interior portions) was digested over night at 56°C in 20 ul proteinase K and 180 ul Buffer ATL from the Qiagen DNeasy Blood & Tissue Kit. The gDNA was extracted from the digest by the Qiagen AllPrep PowerFecal DNA/RNA Kit following manufacturers protocol (Qiagen 80244).

### Quality assessment and library preparation

Species identification was confirmed for SM samples as the population is sympatric with Sitka black-tailed deer (*Odocoileus hemionus sitkensis*). The mitochondrial D-Loop was amplified using L15527 and H00438 primers (Wu et al. 2003) and sequenced on an ABI 3730 (Applied Biosystems). The consensus sequences generated in BioEdit (v 7.0.4.1) were screened using NCBI BLAST to identify the species. For all known mountain goat fecal samples DNA concentrations were measured with a Qubit 3.0 Fluorometer per manufacturers protocol (Invitrogen). Samples were concentrated if the extracted concentration of extracted fecal DNA was below 3 ng/ul. The validated Illumina 16S rRNA Metagenomic Sequencing Library Preparation (#15044223 rev. B) protocol was then followed for library preparation (Rimoldi et al., 2018). The 16S ribosomal ribonucleic acid (16S rRNA) hypervariable region, specifically the V3 and V4 regions, were targeted with four variants of 341F and 805R primers using the primers designed by Herlemann et al., 2011. A unique combination of Nextera XT indexes, index 1 (i7) and index 2 (i5) adapters were assigned to each sample for multiplexing and pooling.

Four replicates of each sample of fecal gDNA were amplified in 25 ul PCR using the 341F and 805R primers. The replicated amplicons for each sample were combined into a single reaction of 100 ul and purified using a QIAquick PCR Purification Kit (Qiagen, 28104) and quantified on the Qubit Fluorometer. Sample indexes were annealed to the amplicons using an 8-cycle PCR reaction to produce fragments approximately 630 bp in length that included ligated adaptors; the target amplicon is approximately 430 bp in length (Illumina 16S rRNA Metagenomic Sequencing Library Preparation; #15044223 rev. B). Aliquots of 100 ng DNA for each sample were pooled together and purified with the QIAquick PCR Purification Kit for a final volume of 50 ul. The final purified library was validated by TapeStation (Agilent, G2991AA) and sequenced in 300 bp pair-end reads on an Illumina MiSeq sequencer at the Genomic Facility of Guelph University (Guelph, Ontario).

## Analysis

### Bioinformatics and taxonomic evaluation

The quality of the raw sequences was assessed with FastQC (v 0.11.4) and the low-quality cut-off for forward and reverse reads was determined. Forward and reverse reads were then imported into QIIME2 (v 2018.6) for quality control, diversity analysis, and sequence classification. The quality control function within QIIME2, DADA2, was used to truncate forward and reverse reads and perform denoising, and the detection and removal of chimeras. The results of DADA2 with only forward, reverse, and merged reads were analyzed independently; note QIIME2 follows the curated DADA2 R library workflow (https://benjjneb.github.io/dada2/) that requires zero mismatches in overlapping reads for successful merging, since reads are denoised and errors are removed before merging occurs. Sequencing data were grouped by status (captive or wild), population (CZ, CR, SM, or YB), and collection time (samples from CR collected in June, July, or August) for analytical purposes.

Alpha diversity estimates of community richness included Shannon Index and observed OTUs and estimates of community evenness included Pielou’s evenness. A phylogenetic tree was developed in QIIME2 for beta diversity estimates (Supplementary Figure S6). Pairwise sample estimates (beta diversity) included Bray-Curtis, unweighted UniFrac, and weighted UniFrac dissimilarity distance matrices. The taxonomy, to the species level, of all sample reads were assigned using Silva 132 reference taxonomy database (https://docs.qiime2.org/2019.1/data-resources/). The relative proportion of *Firmicutes* to *Bacteroidetes* was calculated for each of the grouped data as variation in the ratio is associated with individual body condition (Donaldson et al., 2006; Ley et al., 2006). For this analysis samples within the groupings were normalized and rarified to the sample with the fewest contigs (6,096 contigs for population and status; and, 7,342 contigs for collection month).

### Statistical analysis and visualization

Differences in community richness and evenness by groupings were assessed with the Kruskal-Wallis rank sum test and p-values were adjusted using Benjamini & Hochberg correction (q-values; Storey, 2002). All taxonomic analysis and visualization were computed with the unclassified reads removed. A Wilcoxon rank sum test with continuity correction was used to assess the differences in relative abundance of taxonomic classifications based on origin with all unclassified sequences removed. A Kruskal-Wallis rank sum test was used to assess the differences in relative abundance of taxonomic classifications based on collection month. Differences were treated as significant if the p-value was < 0.01 and a post-hoc (Dunn) test was conducted to determine where the differences occurred between the three collection times.

Statistical PERMANOVA tests were conducted using the ADONIS function from the R package Vegan (v 2.5.2) on Bray Curtis dissimilarity matrices to test for the presence of shifts in the microbiome communities between groups. A detrended correspondence analysis (DCA) was conducted to determine significant communities between groups at each taxonomic level, with only the taxonomic level with the highest resolution (species) reported and visualized. A principle coordinates analysis (PCoA) was also conducted using Bray Curtis dissimilarity matrices. Data were visualized with the ggplot2 (v 2.3.0.0) package in RStudio (v 3.4.1). Additional materials associated with the analysis are available on Dryad (doi:10.5061/dryad.kk17v5d) and relevant scripts are available on GitLab (https://gitlab.com/WiDGeT_TrentU).

## Results

### Samples and quality filtering

A total of 54 fecal samples were selected for this study, but seven samples from SM were excluded because of > 95% identification as black-tailed Sitka deer and five were excluded for quality reasons. A total of ~ 5.37 million paired-end reads were generated from the remaining 42 fecal samples (SRA accession number PRJNA522005). FastQC analysis indicated that both forward and reverse reads lost quality > 250 bp in length (Phred score < 25), so all reads were trimmed to a length of 250 bp and following DADA2 quality filtering, resulted in 896,534 high quality overlapping reads (contigs) kept for taxonomic and diversity analyses. The sequence breakdown for each group can be seen in Supplementary Table S2. Analyses were also conducted on each read set individually, but we only report the results for the merged reads as the patterns were qualitatively similar (just considerably more data). Contigs that were classified to Archaeal lineages (6,013 contigs) were removed from the analysis. The remaining 890,521 contigs were classified into 3,886 unique 16S rRNA ribosomal sequence variants (RSVs) with at least 1 representative sequence, and 3,854 unique RSVs with at least 10 representative sequences.

### Assessing diversity and species composition across groups

The three alpha diversity metrics did not show any differences between the captive and wild populations (q-value > 0.89 for all comparisons; Supplementary Table S3; Figure 2); however, there were more unique classifications in wild than in captive mountain goats (Supplementary Table S4). Across the four populations, the alpha diversity metrics did not differ (q-value > 0.57 for all comparisons), but moderate differences between the collection months at CR were observed (q-value = 0.04 for all comparisons; Supplementary Table S3; Figure 2), with a trend toward decreasing diversity as the summer progressed. The significant differences (q-value < 0.01) in the taxonomies observed between June-July, June-August, and July-August at CR were contributed by 3 classes, 4 families, 3 genera, and 3 species (Supplementary Table S6), and there was no difference in the phyla or orders observed between any months.

**Figure 2.**
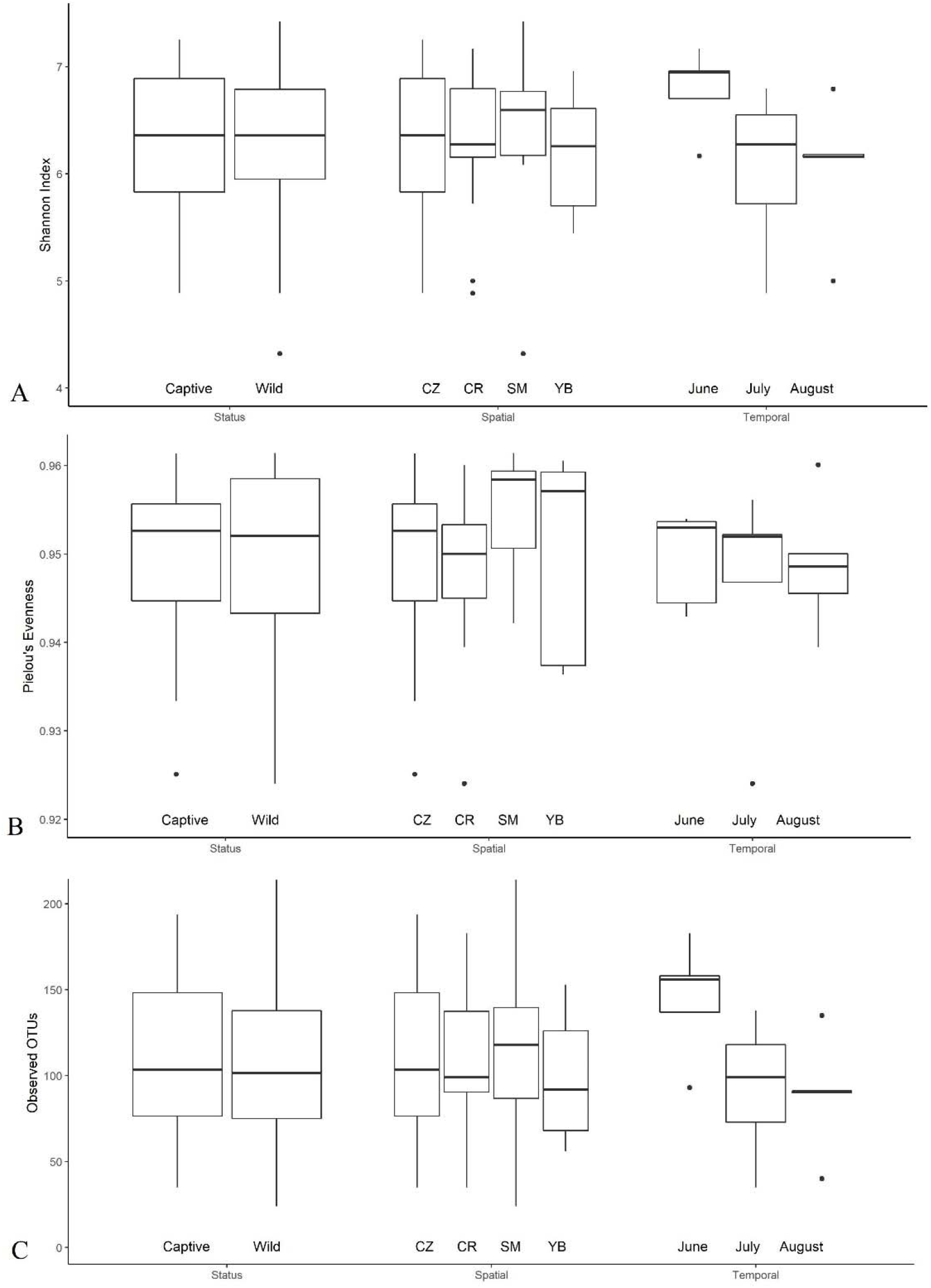
Median (horizontal lines), range (vertical lines) and interquartile range (box) from calculated alpha diversity measured from rarified samples for captive and wild mountain goats, spatial and temporal comparisons: (A) Shannon Index reflecting community diversity, (B) Pielou’s community evenness and (C) observed number of OTUs.

The top two phyla based on averaged relative abundance were *Firmicutes* and *Bacteroidetes* (Table 1; Table 2). The important distinguishing taxonomic differences (p-value < 0.01) between captive and wild mountain goats arose from a suite of different groups, ranging from three different phyla to 26 different species (Supplementary Table S5; Figure 3). Notably of the top five groups, they only differed by a *Spirochaetes* identification for captive and by *Proteobacteria* for wild mountain goats (Table 1; Table 2). A similar pattern was observed across populations (Table 1; Table 2) and likewise, *Firmicutes* and *Bacteroidetes* ratios were more-or-less consistent, and only one genus differed across months (Table 2).

**Table 1.**
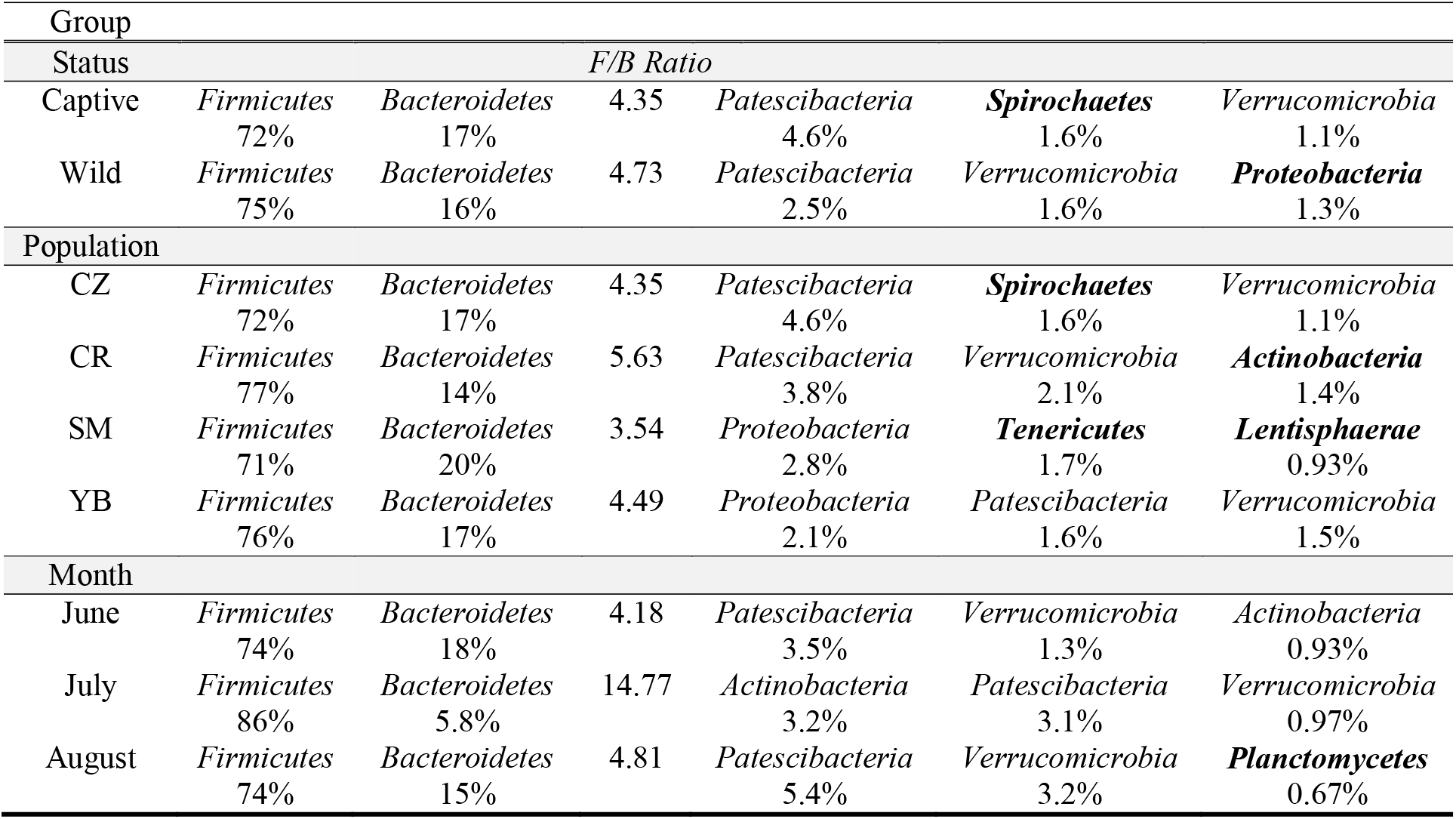
The top five microbiome phyla based on relative abundance for the four mountain goat populations (Calgary Zoo, Alberta = CZ; Caw Ridge, Alberta = CR; Ship Mountain, Alaska = SM; and Yes Bay, Alaska = YB), for captive and wild mountain goats, and for the different collection months at CR. The relative *Firmicutes* to *Bacteroidetes* ratios are also reported. Bolded groups represent a unique classification among groups.

**Table 2.**
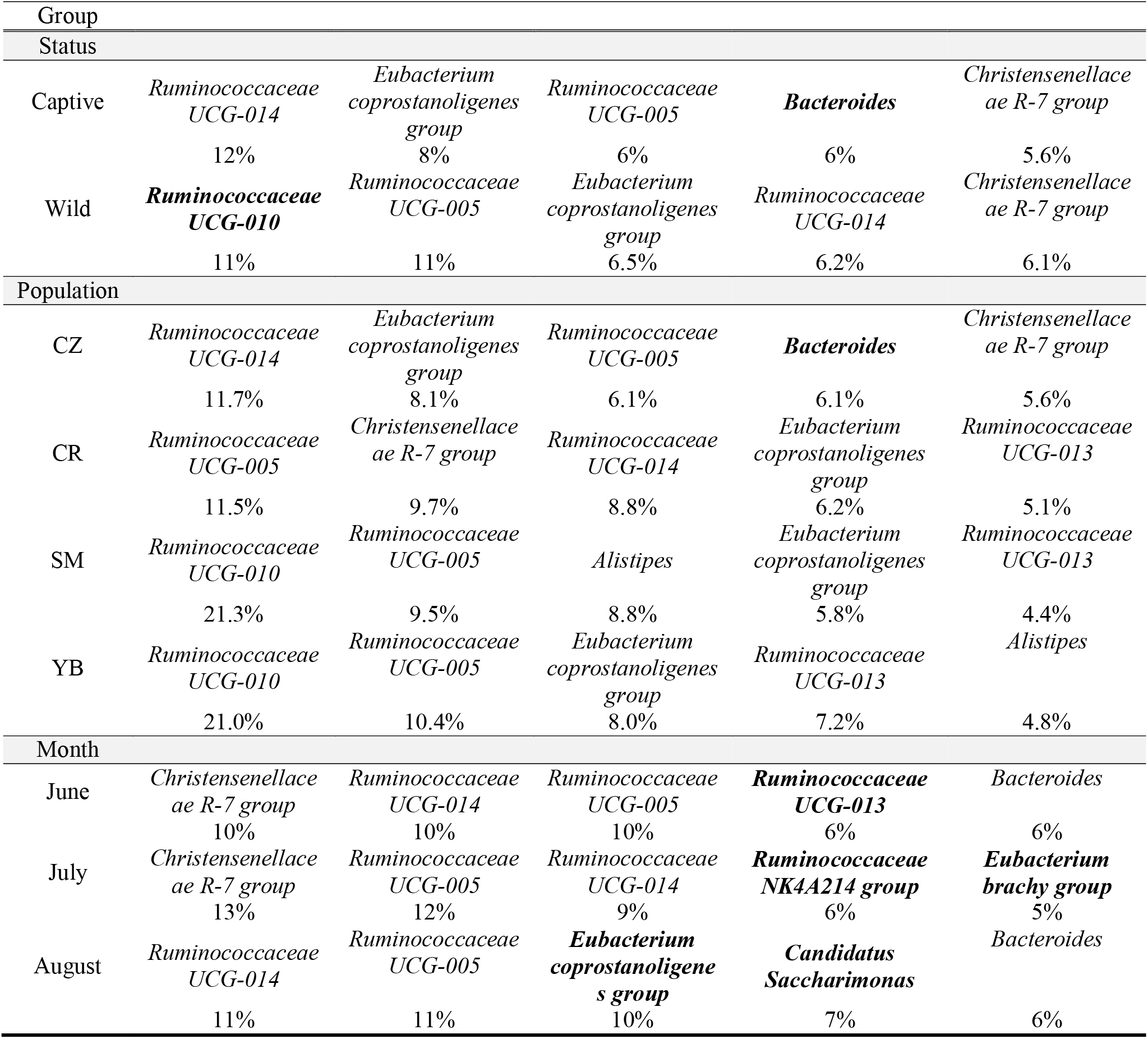
The top five microbiome genera, with relative percent classification, for all four mountain goat populations, for captive and wild mountain goats, and for the different collection months at Caw Ridge, Alberta. Bolded groups represent a unique classification among groups.

**Figure 3.**
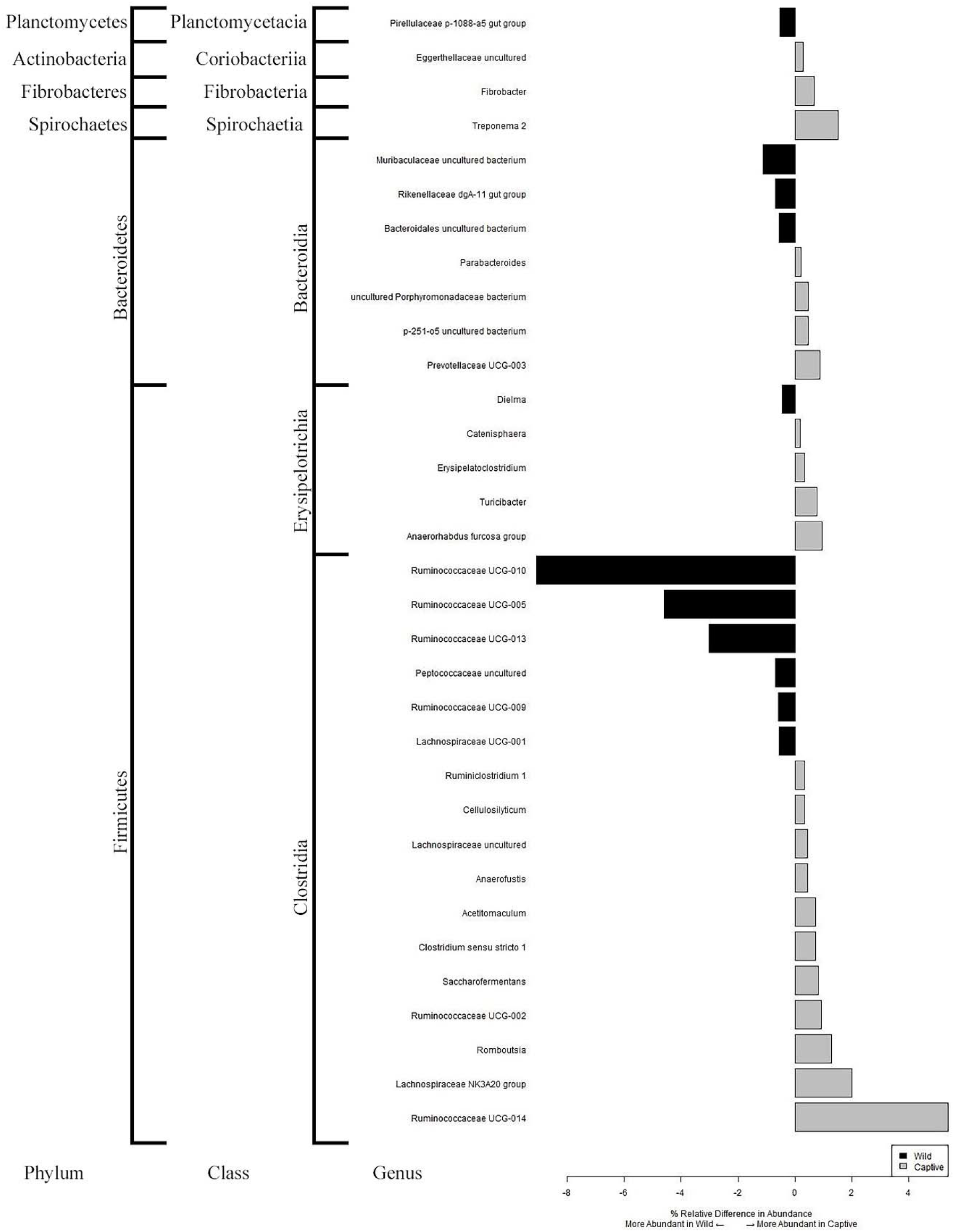
The difference in relative percentage abundance of statistically significant genera (P-value <0.01) that differ between captive and wild populations of mountain goats and their taxonomic breakdown at the phylum and class levels.

### Quantifying drivers of microbial diversity

A 4-way PERMANOVA of the Bray Curtis dissimilarity matrix indicated that the captive and wild mountain goats had significant shifts (p<0.001) in their microbiome communities (Table 3; Figure 2). The same pattern was observed across populations (p<0.001) and over time (p=0.018; Table 3). The DCA (at the species level) clearly separated captive from wild as well as the four sampled populations, with the percent variance explained (eigenvalues) being 46.00% and 24.37% for DCA1 and DCA2, respectively (Figure 4). Similarly, the PCoA at the species level, based on Bray Curtis dissimilarity, weighted UniFrac, and unweighted UniFrac, clearly separated captive from wild as well the four sampled populations (Supplementary Figure S4-S6). No temporal groupings were observed (Figure 4; Supplementary Figures S4-S6), despite the PERMANOVA patterns (Table 3).

**Table 3.**
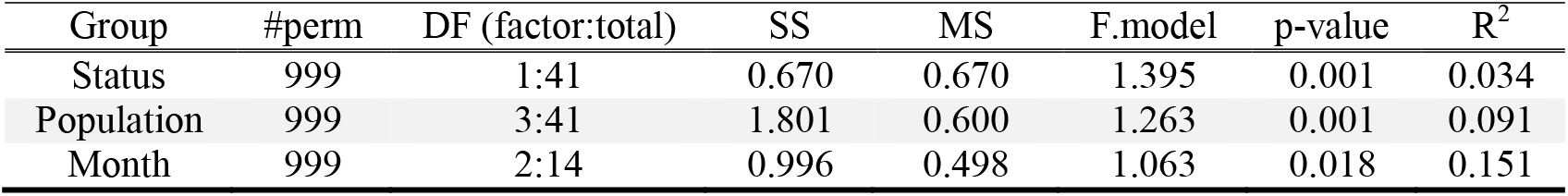
Results of 4-way, 2-way, and 3-way PERMANOVA comparisons of Bray Curtis dissimilarity matrix for all four mountain goat populations, for captive and wild mountain goats, and for the different collection months at Caw Ridge, Alberta, respectively.

| Group | #perm | DF (factor:total) | SS | MS | F.model | p-value | R<sup>2</sup> |
| --- | --- | --- | --- | --- | --- | --- | --- |
| Status | 999 | 1:41 | 0.670 | 0.670 | 1.395 | 0.001 | 0.034 |
| Population | 999 | 3:41 | 1.801 | 0.600 | 1.263 | 0.001 | 0.091 |
| Month | 999 | 2:14 | 0.996 | 0.498 | 1.063 | 0.018 | 0.151 |

**Figure 4.**
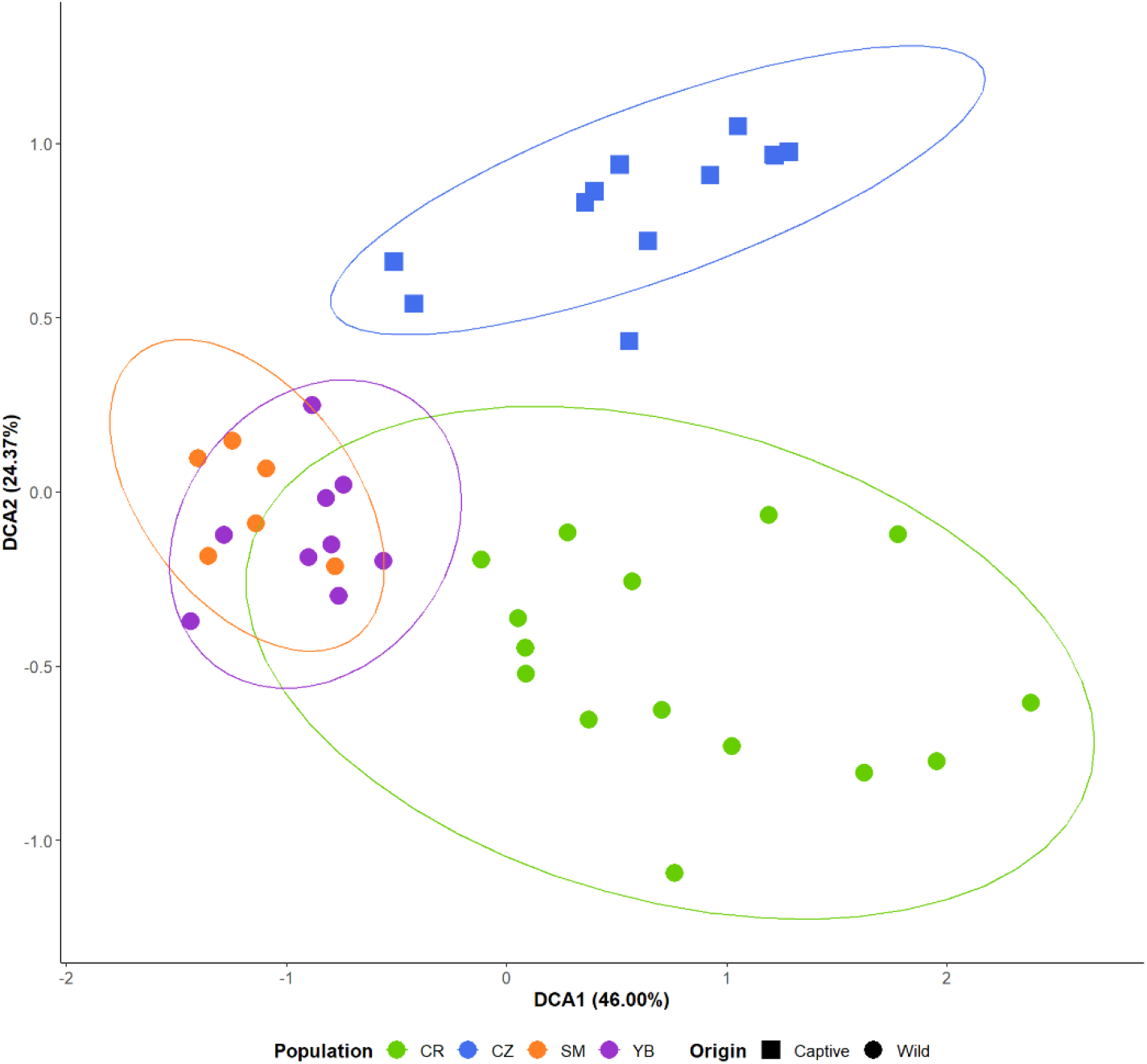
Visualization of the DCA conducted from the species level classification counts by origin and status. The distinction between origin, either Alberta (CR and CZ samples) or Alaska (SM and YB samples), is made clear by DCA1 which explains 46.00% of the variance. The distinction between captive (circle samples) and wild (triangle samples) populations are made clear by DCA2 which explains 26.37% of the variance.

## Discussion

The structure of the fecal microbiome is influenced by a multitude of biotic and abiotic host-specific factors, including genetics, diet, environment, and health status (Bahrndorff et al., 2016). While unaccounted factors such as age and sex likely contribute to variation in the microbiome (Kook et al., 2018; Bennett et al., 2016; Million et al., 2013), we observed clear effects of captivity (Fig. 3; 4) and trends suggestive of a seasonal effect (Fig. 2; Table 3. This study should therefore inform both basic and applied research of ungulate microbiomes, and potentially inform the management of this enigmatic species by identifying the composition of microbial populations in wild, healthy individuals. Specifically, there is a mounting body of evidence that links the fecal microbiome to the health of individuals and these links may be useful tools in guiding population management in captive and wild populations (Bahrndorff et al., 2016; Pannoni, 2015). Moreover, given the links between host-microbiome and adaptive responses (Webster et al., 2017; Ziegler et al. 2017), it is conceivable that population-level microbiome divergence could, and likely should, be factored into the designation of evolutionary significant units (*sensu* Moritz, 1994) and predictive models as it pertains to adaptive responses.

### Comparing diversity metrics

Most mammalian microbiome comparisons to date have shown significant decreases in the alpha diversity (diversity and evenness) in captive ruminant populations; however, there are also studies that show no changes or even increases in diversity in captive populations in other mammals (McKenzie et al., 2017; Kohl et al., 2014). In Bovidae, McKenzie et al. (2017) showed no statistical difference between captive and wild groups, which was also observed in this study on mountain goats. This finding suggests that, for the mountain goats at the Calgary Zoo, captivity has played little effect on overall microbiome diversity, but rather microbiome community structure (Figures 2, 3). Community diversity estimates for mountain goats were also comparable to other unique ruminant populations, such as muskoxen (*Ovibos moschatus*), Bactrian camel (*Camelus bactrianus*), and Norwegian reindeer (*Rangifer tarandus tarandus*), but less comparable to roe deer (*Capreolus capreolu*) and dzo (*Bos grunniens* and *Bos taurus* hybrid; He et al., 2018; Salgado-Flores et al., 2016).

There are strong seasonal dynamics in activity, diet, and body condition in wild mountain goats that could lead to disparity between the microbiome community diversity we measured in mountain goats, and other studies from ruminant species. In particular, mountain goats appear to lose and regain a significant amount of body mass (up to 30-40%) between winter and summer (Côté and Festa-Bianchet, 2003) and are exposed to environmental extremes (e.g. 0 to 25°C). Such dynamics may provide important context for understanding seasonal changes in the microbiome. For example, during spring and early summer animals, particularly parturient females, might be oriented to obtaining, absorbing, and utilizing nutrients whereas later in late-summer and fall may be storing nutrients for periods of winter scarcity.

The dominance of *Firmicutes* and *Bacteroidetes* phyla is consistent with other ruminant studies (O’ Donnell et al., 2017). There were minimal changes in the relative percent *Firmicutes* (from 75% to 72%) and increases in the relative classified *Bacteroidetes* (from 16% to 17%) from wild to captive mountain goats, which is consistent with the general mammalian trends observed in McKenzie et al. (2017). The *Firmicutes* to *Bacteroidetes* ratio (F/B) has been used to give a rough estimate of the function of the microbiome, where a higher F/B ratio in treatment groups compared to control groups in mice and humans has generally been linked with diseases such as obesity and an elevated body mass index (BMI; Koliada et al., 2017). Koliada et al., (2017) suggested that the association between increase F/B ratios and elevated BMIs arises from *Firmicutes*, a group linked with nutrition absorption and circulation, being more efficiently able to participate in energy utilization than *Bacteroidetes*, a group more associated with nutrition storage. Therefore, increases in the relative abundance of *Firmicutes* can significantly contribute to the hosts’ elevated BMI phenotype. The diets of ruminants are typically low in nutrients, especially during winter months, and the elevated F/B ratio in mountain goats detected in our summer samples, relative to some other ruminants, suggests that the metabolism efforts of mountain goat microbiomes are driven to obtain and absorb, rather than store, nutrition; a pattern consistent with expected seasonal energy balance strategies of northern ungulates. *Actinobacteria*, another phylum detected in mountain goats, is associated with body condition (e.g. BMI) and might contribute to the immune system of the host (Koliada et al., 2017; Ventura et al., 2007). An important next step will be to link F/B ratio and relative abundance of *Actinobacteria* to phenotypic attributes like body condition and mass in mountain goats, which is possible in both a captive (CZ) and wild (CR) setting.

### Detectable shifts in microbiome community compositions

There were significant shifts in the fecal microbiome community composition between the four different population groups. However, an R^2^ value of 0.09 suggests that there are multiple factors, beyond population origin, driving the shifts seen in the fecal microbiome communities. Unmeasured factors like age, sex, and reproductive status would likely explain some of the remaining variation; still, there was considerably less variation explained by captive and wild group designation (R^2^=0.03). The spatial differences thus are more prominent in shaping fecal microbial composition than that of captivity, and our model fit is consistent with other comparisons of captive and wild groups of ruminants (McKenzie et al., 2017). Here we speculate that the northern latitude or geographic isolation of the Alaskan population has contributed to reduced fecal microbiome diversity in terms of species number (Shafer et al., 2012; Table S4), while the shifts at the phyla level are consistent with diet alterations between captive and wild animals with similar patterns seen in Sika deer (Guan et al., 2017).

Interestingly, sample time at CR had the best model fit (R^2^=0.15; p=0.018), where the most different genera (p < 0.001) between June and July was the *Eubacterium hallii group* (increased in July); similarly, *Clostridiales family XIII AD3011 group* and *Eubacterium hallii groups* increased in July. The *Eubacterium hallii group* is a commonly observed genus in mammal microbiomes and plays a role in glycolysis (Engels et al., 2016); we suggest that the increase of *Eubacterium hallii group* in July relative to June or August might be associated with shifts in forage quality during the peak of summer at CR. Importantly, while the core fecal microbiome appears relatively constant in the mountain goat, at the local-level there are clear shifts in individual bacteria that reflect changes in the microhabitat over time.

## Conclusion

We are beginning to understand how the fecal microbiome influences the host and is, in turn, influenced by the host. There is a need to understand how to best apply this knowledge to aid the management and conservation of mammal populations (Li et al., 2018; O’ Donnell et al., 2017). The direct connection of the fecal microbiome to both individual and population health makes fecal microbiome assays an important tool for monitoring the health and disease trends of domesticated, captive, and wild populations of mammals (Bahrndorff et al., 2016; Jiménez & Sommer, 2017). This study shows that there are clear differences in fecal microbiome community composition, but not diversity, that can be best explained by a combination of factors, including status, seasonality and population of origin. The baseline microbiome data described here thus has the potential to provide valuable insight into the health of wild mountain goat populations and represents an important frame of reference for the development of future monitoring programs and associated management strategies.

## Acknowledgments

We thank Doug Whiteside (Calgary Zoo), Daria Martchenko, Boyd Porter, and Yasaman Shakeri for helping with sample collection. Jess Breen and Spencer Anderson helped with aspects of library preparation. Justin Johnson helped develop the sample map. Erica Newton for support in R statistical analysis and figure development. We thank three anonymous reviewers for their comments that improved this manuscript. The authors declare that they have no competing interests.

## References

Amato KR, Sander JG, Song SJ et al. Evolutionary trends in host physiology outweigh dietary niche in structuring primate gut microbiomes. ISME J 2018, DOI: 10.1038/s41396-018-0175-0.

Amato KR, Yeoman CJ, Kent A et al. Habitat degradation impacts black howler monkey (*Alouatta pigra*) gastrointestinal microbiomes. ISME J 2013;7:1344–53.

Bahrndorff S, Alemu T, Alemneh T et al. The microbiome of animals: implications for conservation biology. Int J Genomics 2016:5304028, DOI:10.1155/2016/5304028.

Bennett G, Malone M, Sauther ML et al. Host age, social group, and habitat type influence the gut microbiota of wild ring-tailed lemurs (*Lemur catta*). Am J Primatol 2016;78:883–92.

Berg RD. The indigenous gastrointestinal microflora. Trends Microbiol 1996;4:227–310.

Bik EM, Costello EK, Switzer AD et al. Marine mammals harbor unique microbiotas shaped by and yet distinct from the sea. Nat Commun 2016;7:10516.

Breen J, Britt M, Martchenko D et al. Extensive field-sampling reveals the uniqueness of a trophy mountain goat population. J Wildlife Manag 2018, DOI: 10.1101/484592.

Carey HV, Walters WA, Knight R. Seasonal restructuring of the ground squirrel gut microbiota over the annual hibernation cycle. Am J Physiol Regul Integr Comp Physiol 2013;304:R33–RR42.

Carthey AJR, Gillings MR, Blumstein DT. The extended genotype: microbially mediated olfactory communication. Trends Ecol and Evol 2018;33:885–94.

Clayton JB, Al-Ghalith GA, Long HT et al. Associations between nutrition, gut microbiome, and health in a novel nonhuman primate model. Sci Rep 2018;8:11159.

Côté SD, Festa-Bianchet M. Mountain goat. In: Feldhamer GA, Thompson BC, Chapman JA (eds). Wild mammals of North America: biology, management and conservation, 2nd edn. The John Hopkins University Press, Baltimore, Md, 2003, 1061–75.

Dikongué E and Ségurel L. Latitude as a co-driver of human gut microbial diversity? Bioessays 2017;39.

Di Marco M, Boitani L, Mallon D et al. A retrospective evaluation of the global decline of carnivores and ungulates. Conserv Biol 2014;28:1109–18.

Donaldson GP, Lee SM, Mazmanian SK. Gut biogeography of the bacterial microbiota. Nat Rev Microbiol 2015;14:20–32.

Engels C, Ruscheweyh HJ, Beerenwinkel N et al. The common gut microbe *Eubacterium hallii* also contributes to intestinal propionate formation. Front Microbiol 2016;7:713.

Feng Q, Chen WD, Wang YD. Gut microbiota: an integral moderator in health and disease. Front Microbiol 2018;9:151.

Festa-Bianchet M, Côté SD. Mountain goats: ecology, behavior, and conservation of an alpine ungulate (1st ed.). Washington, D.C.: Island Press, 2008. Festa-Bianchet M. Oreamnos americanus. The IUCN red list of threatened species. 2008 DOI: 10.2305/IUCN.UK.2008.RLTS.T42680A10727959.en.go

Fox JL, Smith CA. Winter mountain goat diets in southeast Alaska. J Wildlife Manag 1988;52:362–5.

Guan Y, Yang H, Han S et al. Comparison of the gut microbiota composition between wild and captive sika deer (Cervus nippon hortulorum) from feces by high-throughput sequencing. AMB Express 2017;7:212.

Hamel S, Côté SD. Habitat use patterns in relation to escape terrain: are alpine ungulate females trading off better foraging sites for safety? Can J Zool 2007;85:933–43.

He J, Yi L, Hai L et al. Characterizing the bacterial microbiota in different gastrointestinal tract segments of the Bactrian camel. Sci Reports 2018;8:654.

Herlemann DP, Labrenz M, Juřgens K et al. Transitions in bacterial communities along the 2000km salinity gradient of the Baltic Sea. ISME J 2011;5:1571–9.

Hicks AL, Lee KJ, Couto-Rodriguez M et al. Gut microbiomes of wild great apes fluctuate seasonally in response to diet. Nat Commun 2018;9:1786.

Huntington G, Woodbury M, Anderson VPAS. Invited review: growth, voluntary intake, and digestion and metabolism of North American bison. Applied Animal Sci 2019;35:146–60.

Hu X, Liu G, Li Y et al. High-throughput analysis reveals seasonal variation of the gut microbiota composition within forest musk deer (Moschus berezovskii). Front Microbiol 2018;9:1674.

Ingala MR, Simmons NB, Wultsch C et al. Comparing microbiome sampling methods in a wild Mammal: fecal and intestinal samples record different signals of host ecology, evolution. Front Microbiol 2018;9:803.

Jiméenez RR, Sommer S. The amphibian microbiome: natural range of variation, pathogenic dysbiosis, and role in conservation. Biodivers Conserv 2017;26:763–86.

Khan MJ, Gerasimidis K, Edwards CA et al. Role of gut microbiota in the aetiology of obesity: proposed mechanisms and review of the literature. J Obesity 2016, DOI: 10.1155/2016/7353642.

Kohl KD, Skopec MM, Dearing MD. Captivity results in disparate loss of gut microbial diversity in closely related hosts. Conserv Physiol 2014;2:cou009.

Koliada A, Syzenko G, Moseiko V et al. Association between body mass index and Firmicutes/Bacteroidetes ratio in an adult Ukrainian population. BMC Microbiol 2017;17:120.

Kook SY, Kim Y, Kang B et al. Characterization of the fecal microbiota differs between age groups in Koreans. Intest Res 2018;16:246–54.

Kreisinger J, Schmiedová L, Petrželková A et al. Fecal microbiota associated with phytohaemagglutinin-induced immune response in nestlings of a passerine bird. Ecol Evol 2018;8:9793–802.

Ley RE, Turnbaugh PJ, Klein S et al. Microbial ecology: human gut microbes associated with obesity. Nature 2006;444:1022–3.

Li Y, Hu X, Yang S et al. Comparative analysis of the gut microbiota composition between captive and wild forest musk deer. Front Microbiol 2017;8:1705.

Li Y, Hu X, Yang S et al. Comparison between the fecal bacterial microbiota of healthy and diarrheic captive musk deer. Front Microbiol 2018;9:300.

Mainguy J, Côté SD, Festa-Bianchet M et al. Father–offspring phenotypic correlations suggest intralocus sexual conflict for a fitness-linked trait in a wild sexually dimorphic mammal. Proc R Soc B Biol Sci 2009;276:4067–75.

Martin TE. Abiotic vs. biotic influences on habitat selection of coexisting species: climate change impacts? Ecology 2001;82:175–88.

McKenzie VJ, Song SJ, Delsuc F et al. The effects of captivity on the mammalian gut microbiome. Integr Comp Biol 2017;57:690–704.

Million M, Lagier JC, Yahav D et al. Gut bacterial microbiota and obesity. Clin Microbiol Infect 2013;19:305–13.

Moritz C. Defining ‘evolutionarily significant units’ for conservation. Trends Ecol Evol 1994;9:373–5.

Muegge BD, Kuczynski J, Knights D et al. Diet drives convergence in gut microbiome functions across mammalian phylogeny and within humans. Science 2011;332:970–4.

Nishida AH, Ochman H. (2017). Rates of gut microbiome divergence in mammals. Mol Ecol 2018;27:1884–97.

Noel SJ, Attwood GT, Rakonjac J et al. Seasonal changes in the digesta-adherent rumen bacterial communities of dairy cattle grazing pasture. PLoS ONE 2017;12:e0173819.

O’ Donnell MM, Harris HMB, Ross RP et al. Core fecal microbiota of domesticated herbivorous ruminant, hindgut fermenters, and monogastric animals. Microbiology open, 2017;6:e00509.

Pannoni SB. Developing Microbial Biomarkers to Non-invasively Assess Health in Wild Elk (Cervus canadensis) Populations, Dissertation. University of Montana - Missoula, 2015, Accessed 2019 November at https://scholarworks.umt.edu/utpp/66/.

Poole KG, Heard DC. Seasonal habitat use and movements of mountain goats, Oreamnos americanus, in east-central British Columbia. Can Field Nat 2003;117:565–76.

Rimoldi S, Terova G, Ascione C et al. Next generation sequencing for gut microbiome characterization in rainbow trout (Oncorhynchus mykiss) fed animal by-product meals as an alternative to fishmeal protein sources. PLoS ONE 2018;13:e0193652.

Salgado-Flores A, Bockwoldt M, Hagen LH et al. First insight into the faecal microbiota of the high Arctic muskoxen (Ovibos moschatus). Microbial Genomics 2016;2:e000066.

Shafer ABA, Northrup JM, White KS et al. Habitat selection predicts genetic relatedness in an alpine ungulate. Ecol 2012;93:1317–29.

Sharpton TJ. Role of the gut microbiome in vertebrate evolution. mSystems 2018;3:e00174–17.

Smith FA, Hammond Ji, Balk MA et al. Exploring the influence of ancient and historic megaherbivore extirpation on the global methane budget. PNAS 2016;113:874–9.

Smith CA, Raedeke KJ. Group size and movements of a dispersed, low density goat population with comments on inbreeding and human impact. Bienn Symp North Wild Sheep and Goat Counc 1982;3:54–67.

Storey JD. A direct approach to false discovery rates. J R Statist Soc B 2002;64:479–98.

Sun B, Wang X, Bernstein S et al. Marked variation between winter and spring gut microbiota in free-ranging Tibetan macaques (*Macaca thibetana*). Sci Rep 2016;6:26035.

Vandeputte D, Falony G, Vieira-Silva S et al. Stool consistency is strongly associated with gut microbiota richness and composition, enterotypes and bacterial growth rates. BMJ Gut 2016;65:57–62.

Ventura M, Canchaya C, Tauch A et al. Genomics of Actinobacteria: tracing the evolutionary history of an ancient phylum. Microbiol Mol Biol Rev 2007;71:495–548.

Webster NS, Reusch TBH. Microbial contributions to the persistence of coral reefs. ISME J 2017;11:2167–74.

Wu CH, Zhang YP, Bunch TD et al. Mitochondrial control region sequence variation within the argali wild sheep (*Ovis ammon*): evolution and conservation relevance. Mammalia 2003;67:109–18.

ZieglerM, Seneca FO, YumLK et al. Bacterial community dynamics are linked to patterns of coral heat tolerance. Nat Commun 2017;8:14213.

