## Supplementary Figures for "Space, time, and captivity: quantifying the factors influencing the fecal microbiome of an alpine ungulate"

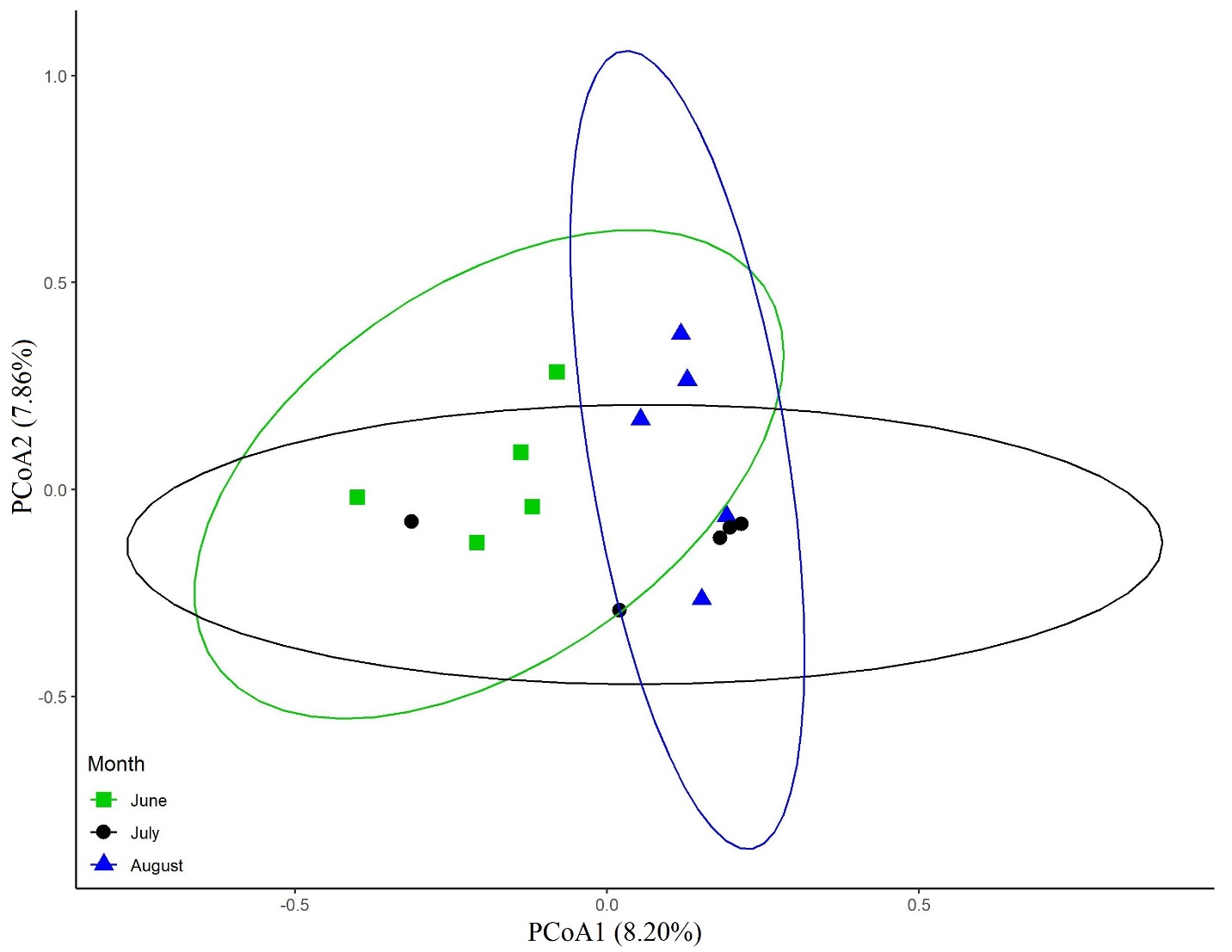

Figure S1. Principal components analysis (PCoA) of Bray Curtis dissimilarity index values. The first component explains 8.2% of the variation and the second component explains 7.9% of the variation. There is no separation of collection months at Caw Ridge.

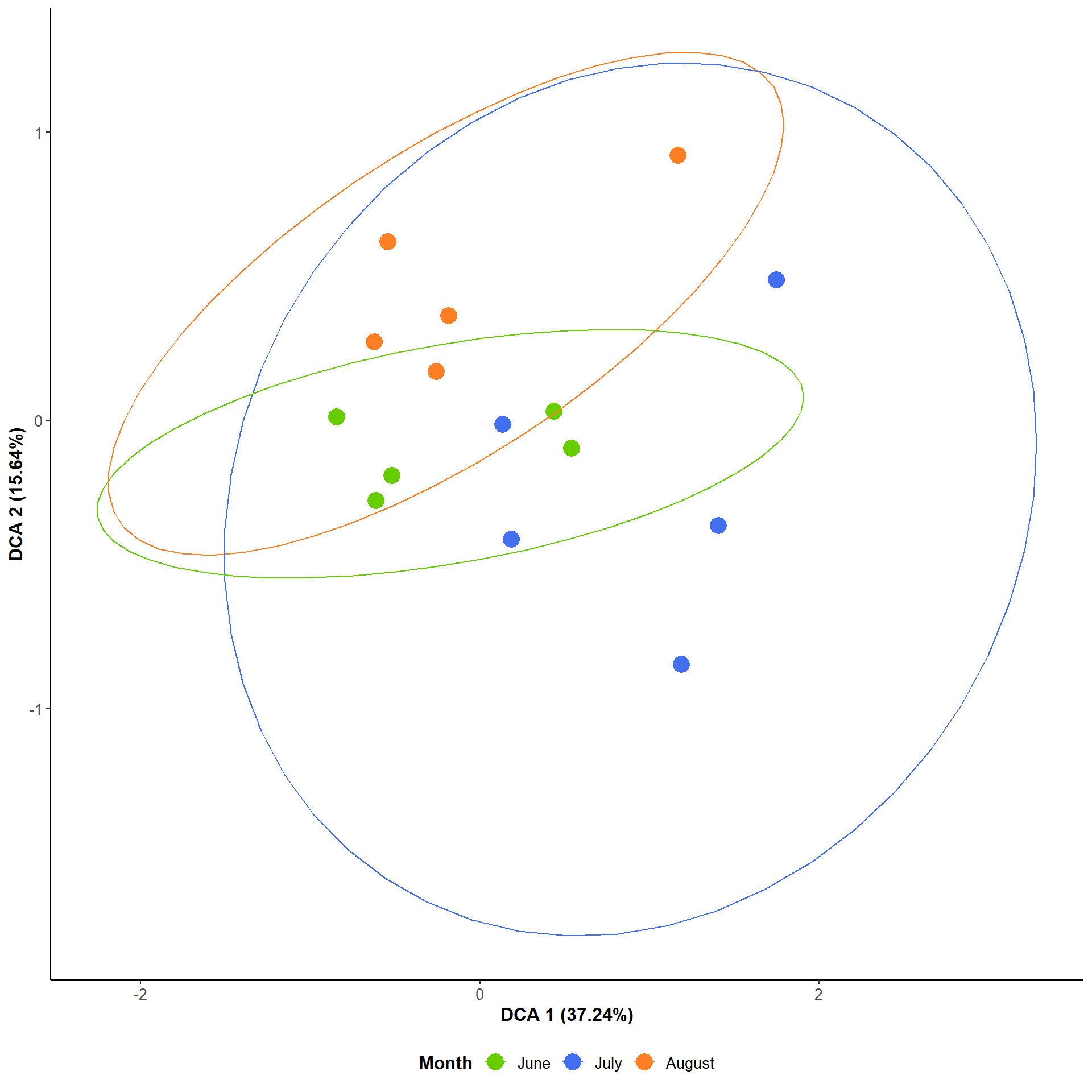

Figure S2. Visualization of the detrended correspondence analysis (DCA) conducted from the species level classification counts, the level with the highest resolution, of the temporally collected Caw Ridge, Alberta samples. The DCA demonstrates the lack of separation by collection month by both DCA1, which explains 37.24% of the variance and by DCA2, which explains 15.64% of the variance.

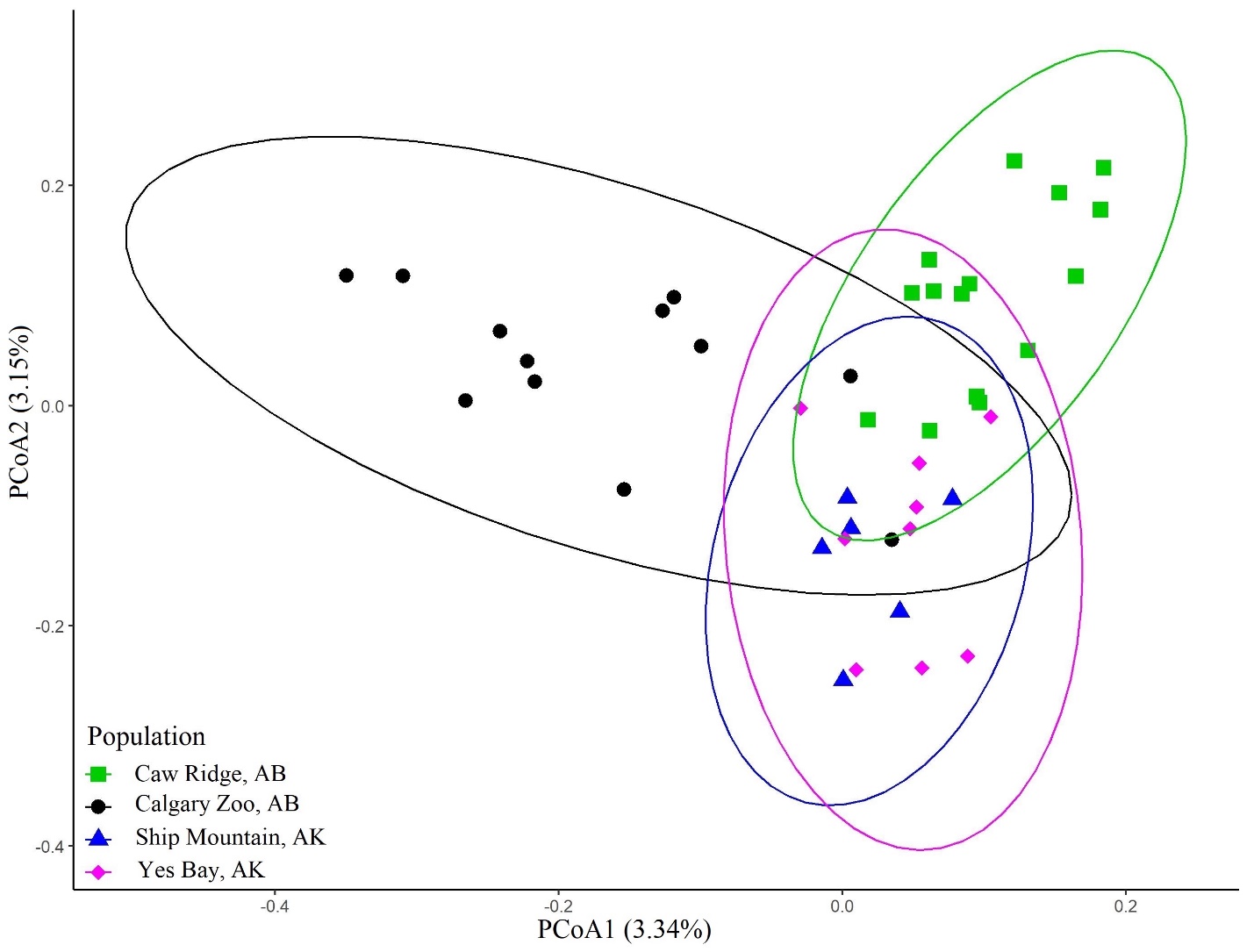

Figure S3. Principal coordinates analysis (PCoA) of Bray Curtis dissimilarity index values at the species level. The first component explains 3.34% of variation and separates the captive population (Calgary Zoo), from the wild populations. The second component explains 3.15% of the variation and moderately separates Caw Ridge, a southern wild population, from both of the northern wild populations.

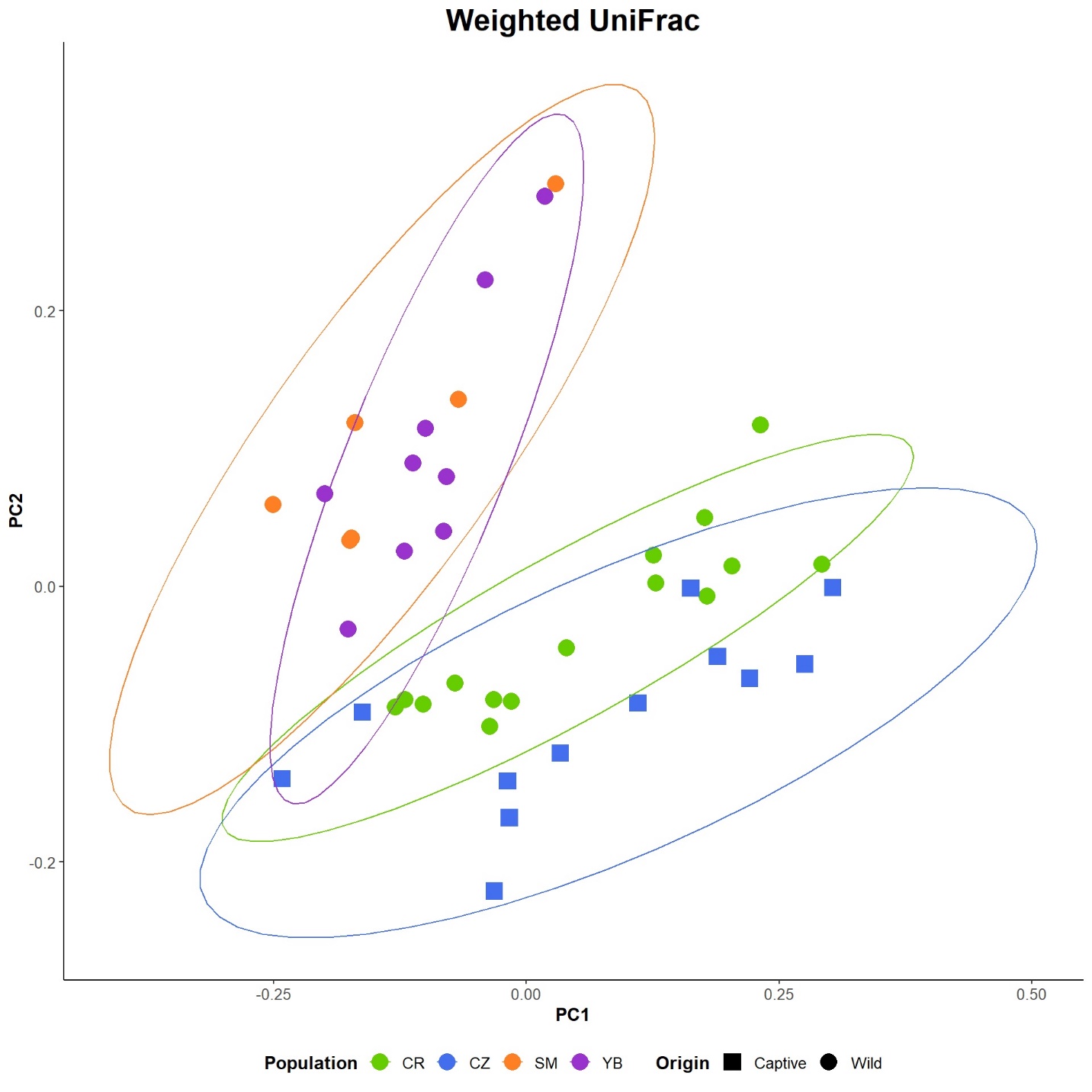

Figure S4. Principal coordinates analysis (PCoA) of weighted UniFrac values at the species level by origin and status. There is a moderate separation between Alberta (CR and CZ samples) and Alaska (SM and YB samples) by PC2, with strong overlap between captive and wild samples.

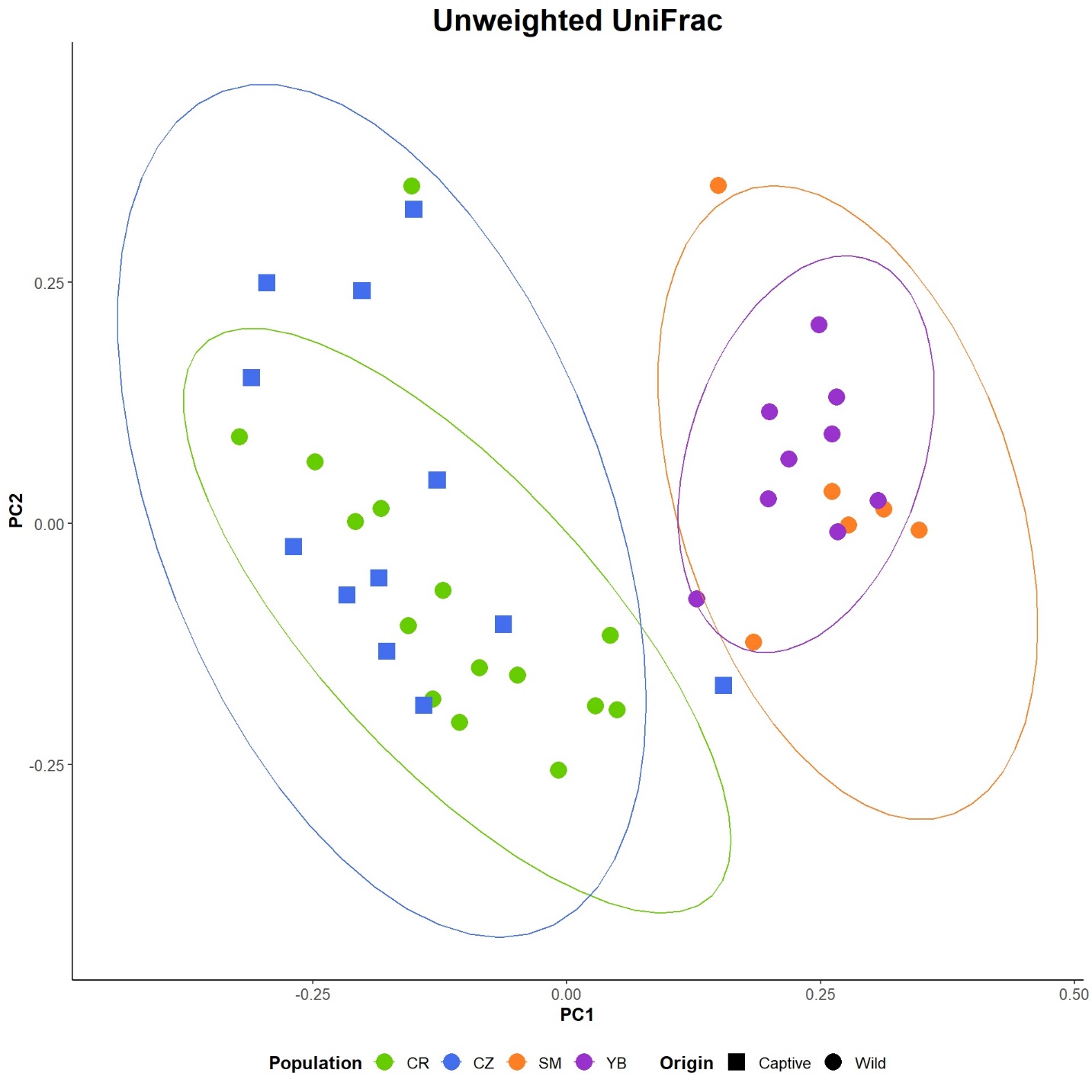

Figure S5. Principal coordinates analysis (PCoA) of unweighted UniFrac values at the species level by origin and status. There is a moderate but clear distinction between Alberta (CR and CZ samples) and Alaska (SM and YB samples) by PC1, with strong overlap between captive and wild samples.
