## Supplementary Tables for "Space, time, and captivity: quantifying the factors influencing the fecal microbiome of an alpine ungulate"

Table S1. Known diet information for the three mountain goat populations (Calgary Zoo, Alberta = CZ; Ship Mountain, Alaska = SM; and Yes Bay, Alaska = YB). No dietary information is available for Caw Ridge, Alberta.

| Group | Diet |
| --- | --- |
| CZ* | Ground beet pulp  RPS canola meal  TM Sul-Pak  Dehydrated alfalfa  Salt  B-Vitamin Pak  Linpro supplement  Limestone 3  ADE vitamin Pak-30, natural  Millrun  Vitamin E acetate 50% ads  Sheep TM micro Lt  Wheat distillers grain 50:50  Dicalcium phosphate 21%  Vitamin D3 blends  Omega 3 milled flax-steel cut  Magnesium oxide 56%  Organic flax oil  Selenium |
| Cleveland Peninsula (SM & YB) | Athyrium spp.  Unknown Fern  Achillea borealis  Caltha leptosepala  Cornus canadensis  Fauria crista-galli  Lupinus nootkatensis  Maianthemum dilatatum  Osmorhiza sp.  Pedicularis spp.  Ranunculus spp.  Saxifraga spp.  Unknown Forb  Valeriana sitchensis  Agrostis spp.  Calamagrostis spp.  Deschampsia spp.  Elymus glacus  Poa spp.  Trisetum spicatum  Unknown Grasses  Lobaria  Peltigera  Unknown Lichen  Sphagnum Moss  Unknown Moss  Carex spp.  Luzula / Juncus  Unknown Sedge/Rush  Rubus spectabilis  Unknown Shrubs  Vaccinium  Thuja plicata |

**Ingredients of herbivore pellets fed ad libidum year-round*

Table S2. Number of sequences included for taxonomic classification for the four mountain goat populations (Calgary Zoo, Alberta = CZ; Caw Ridge, Alberta = CR; Ship Mountain, Alaska = SM; and Yes Bay, Alaska = YB), for captive and wild mountain goats, and for the different collection months at CR.

| Group | Number of sequences |
| --- | --- |
| Status |  |
| Captive | 258,961 |
| Wild | 637,583 |
| Population |  |
| CZ | 258,961 |
| CR | 336,433 |
| SM | 128,250 |
| YB | 172,900 |
| Month |  |
| June | 140,448 |
| July | 98,956 |
| August | 95,929 |

Table S3. Alpha diversity analysis for the four mountain goat populations (Calgary Zoo, Alberta = CZ; Caw Ridge, Alberta = CR; Ship Mountain, Alaska = SM; and Yes Bay, Alaska = YB), for captive and wild mountain goats, and for the different collection months at CR. Alpha diversity metrics include observed OTUs, Shannon Index (community diversity), and Pielou’s evenness (community evenness). All corrected p-values (q-values) were > 0.89 for all comparisons.

| Group | *N* | Mean OTUs | Shannon Index | Pielou’s Evenness |
| --- | --- | --- | --- | --- |
| Status |  |  |  |  |
| Captive | 12 | 111.9±53.4 | 6.28±0.82 | 0.95± 0.01 |
| Wild | 30 | 107.3±44.1 | 6.27±0.72 | 0.95± 0.01 |
| Population |  |  |  |  |
| CZ | 12 | 111.9±53.4 | 6.27± 0.82 | 0.95± 0.01 |
| CR | 15 | 108.7±41.3 | 6.29±1.06 | 0.95± 0.01 |
| SM | 6 | 115.7±57.7 | 6.30±0.68 | 0.95± 0.01 |
| YB | 9 | 99.2±36.4 | 6.22±0.59 | 0.95± 0.01 |
| Month |  |  |  |  |
| June | 5 | 142.6±27.0 | 6.79±0.35 | 0.95± 0.01 |
| July | 5 | 93.4±36.8 | 6.06±0.76 | 0.95± 0.01 |
| August | 5 | 89.8±30.4 | 6.06±0.65 | 0.95± 0.01 |

Table S4. Taxonomic classification breakdown for the four mountain goat populations (Calgary Zoo, Alberta = CZ; Caw Ridge, Alberta = CR; Ship Mountain, Alaska = SM; and Yes Bay, Alaska = YB), for captive and wild mountain goats, and for the different collection months at CR.

| Group | *Phyla* | *Class* | *Order* | *Family* | *Genus* | *Species* |
| --- | --- | --- | --- | --- | --- | --- |
| Status |  |  |  |  |  |  |
| Captive | 14 | 20 | 28 | 55 | 119 | 119 |
| Wild | 15 | 21 | 36 | 64 | 138 | 140 |
| Population |  |  |  |  |  |  |
| CZ | 14 | 20 | 27 | 54 | 118 | 119 |
| CR | 14 | 20 | 28 | 47 | 104 | 109 |
| SM | 13 | 19 | 29 | 45 | 80 | 70 |
| YB | 13 | 19 | 26 | 45 | 86 | 75 |
| Month |  |  |  |  |  |  |
| June | 12 | 18 | 22 | 36 | 78 | 82 |
| July | 13 | 17 | 19 | 30 | 68 | 65 |
| August | 12 | 16 | 19 | 29 | 64 | 65 |

Table S5. Significant results (p<0.01) of the Wilcoxon test comparing relative abundance of taxonomic classifications between captive and wild samples from a total of 3/15 phylum, 4/21 class, 4/36 order, 14/64 family, 34/138 genera, and 25/140 species classifications. The difference in percent relative abundance is shown (relative abundance of captive–relative abundance of wild).

| Level | Classification | p-value | % Diff Rel Abun |
| --- | --- | --- | --- |
| Phylum | Fibrobacteres | 1.20e-6 | 0.58 |
| Phylum | Spirochaetes | 2.99e-6 | 1.32 |
| Phylum | Planctomycetes | 5.95e-6 | -0.44 |
| Class | Fibrobacteria | 1.20e-6 | 0.58 |
| Class | Spirochaetia | 2.99e-6 | 1.32 |
| Class | Planctomycetacia | 5.95e-6 | -0.44 |
| Class | Erysipelotrichia | 1.29e-3 | 1.86 |
| Order | Fibrobacterales | 1.2e-6 | 0.58 |
| Order | Spirochaetales | 2.99e-4 | 1.32 |
| Order | Pirellulales | 5.95e-3 | -0.44 |
| Order | Erysipelotrichales | 1.29e-3 | 1.85 |
| Family | Fibrobacteraceae | 1.20e-6 | 0.58 |
| Family | Eubacteriaceae | 7.40e-6 | 0.39 |
| Family | Clostridiaceae 1 | 1.94e-5 | 0.64 |
| Family | Bacteroidales RF16 group | 1.97e-5 | 0.79 |
| Family | Bacteroidales p-251-o5 | 4.23e-5 | 0.52 |
| Family | Peptostreptococcaceae | 4.56e-5 | 2.35 |
| Family | Tannerellaceae | 2.27e-4 | 0.18 |
| Family | Spirochaetaceae | 2.99e-4 | 1.34 |
| Family | Pirellulaceae | 6.60e-4 | -0.45 |
| Family | Erysipelotrichaceae | 1.29e-3 | 1.88 |
| Family | Bacteroidales uncultured | 2.90e-3 | -1.01 |
| Family | Peptococcaceae | 5.92e-3 | -0.58 |
| Family | Muribaculaceae | 7.08e-3 | -0.95 |
| Family | Clostridiales vadinBB60 group | 7.37e-3 | -1.26 |
| Genus | Anaerorhabdus furcosa group | 9.00e-7 | 0.94 |
| Genus | Eggerthellaceae uncultured | 1.20e-6 | 0.28 |
| Genus | Fibrobacter | 1.20e-6 | 0.66 |
| Genus | Saccharofermentans | 1.20e-6 | 0.81 |
| Genus | Anaerofustis | 7.40e-6 | 0.44 |
| Genus | Prevotellaceae UCG-003 | 1.28e-5 | 0.88 |
| Genus | Clostridium sensu stricto 1 | 1.67e-5 | 0.72 |
| Genus | Turicibacter | 3.41e-5 | 0.76 |
| Genus | Bacteroidales RF16 group uncultured Porphyromonadaceae bacterium | 4.23e-5 | 0.47 |
| Genus | Lachnospiraceae NK3A20 group | 9.01e-5 | 1.98 |
| Genus | Romboutsia | 1.75e-4 | 1.27 |
| Genus | Parabacteroides | 2.27e-4 | 0.20 |
| Genus | Treponema 2 | 2.99e-4 | 1.50 |
| Genus | Pirellulaceae p-1088-a5 gut group | 5.95e-4 | -0.53 |
| Genus | Ruminococcaceae UCG-014 | 7.14e-4 | 5.37 |
| Genus | Acetitomaculum | 1.14e-3 | 0.71 |
| Genus | p-251-o5 uncultured bacterium | 1.14e-3 | 0.47 |
| Genus | Catenisphaera | 1.14e-3 | 0.19 |
| Genus | Lachnospiraceae uncultured | 1.14e-3 | 0.42 |
| Genus | Erysipelatoclostridium | 1.23e-3 | 0.33 |
| Genus | Ruminococcaceae UCG-002 | 1.71e-3 | 0.92 |
| Genus | Ruminococcaceae UCG-005 | 2.30e-3 | -4.59 |
| Genus | Bacteroidales uncultured bacterium | 3.64e-3 | -0.563 |
| Genus | Muribaculaceae uncultured bacterium | 3.87e-3 | -1.13 |
| Genus | Ruminiclostridium 1 | 5.53e-3 | 0.33 |
| Genus | Dielma | 5.59e-3 | -0.46 |
| Genus | Lachnospiraceae UCG-001 | 5.59e-3 | -0.57 |
| Genus | Ruminococcaceae UCG-013 | 6.09e-3 | -3.02 |
| Genus | Cellulosilyticum | 6.09e-3 | 0.35 |
| Genus | Peptococcaceae uncultured | 7.66e-3 | -0.70 |
| Genus | Ruminococcaceae UCG-010 | 8.62e-3 | -9.07 |
| Genus | Rikenellaceae dgA-11 gut group | 8.77e-3 | -0.70 |
| Genus | Ruminococcaceae UCG-009 | 9.91e-3 | -0.60 |
| Species | Anaerorhabdus furcosa group uncultured rumen bacterium | 1.2e-6 | 1.53 |
| Species | Eggerthellaceae uncultured bacterium | 1.2e-6 | 0.55 |
| Species | Fibrobacter uncultured bacterium | 1.2e-6 | 1.28 |
| Species | Anaerofustis uncultured bacterium | 7.4e-6 | 0.76 |
| Species | Prevotellaceae UCG-003 uncultured Bacteroidales bacterium | 1.28e-6 | 1.46 |
| Species | Turicibacter uncultured bacterium | 3.41e-5 | 1.47 |
| Species | Bacteroidales RF16 group uncultured Porphyromonadaceae bacterium | 4.23e-5 | 0.91 |
| Species | Treponema 2 uncultured bacterium | 1.67e-4 | 3.03 |
| Species | Ruminococcaceae UCG-014 uncultured bacterium | 2.24e-4 | 4.39 |
| Species | Family XIII AD3011 group uncultured bacterium | 3.36e-4 | 1.98 |
| Species | Ruminococcaceae UCG-013 uncultured bacterium | 7.61e-4 | -5.08 |
| Species | Ruminococcaceae UCG-014 unidentified rumen bacterium JW32 | 8.68e-4 | 1.23 |
| Species | Erysipelatoclostridium uncultured bacterium | 9.76e-4 | 0.67 |
| Species | p-1088-a5 gut group uncultured bacterium | 9.99e-4 | -0.76 |
| Species | Acetitomaculum uncultured rumen bacterium | 1.14e-3 | 0.73 |
| Species | p-251-o5 uncultured bacterium | 1.14e-3 | 0.91 |
| Species | Catenisphaera uncultured rumen bacterium | 1.14e-3 | 0.37 |
| Species | Defluviitaleaceae UCG-011 uncultured rumen bacterium | 1.14e-3 | 0.26 |
| Species | Ruminococcaceae UCG-002 uncultured bacterium | 1.71e-3 | 1.81 |
| Species | Ruminococcaceae UCG-010 uncultured bacterium | 2.04e-3 | -7.80 |
| Species | Bacteroidales uncultured bacterium | 3.64e-3 | -1.01 |
| Species | Muribaculaceae uncultured bacterium | 4.23e-3 | -1.94 |
| Species | Ruminococcus flavefaciens | 5.53e-3 | 0.26 |
| Species | Dielma uncultured bacterium | 5.58e-3 | -0.83 |
| Species | Rikenellaceae dgA-11 gut group uncultured bacterium | 7.40e-3 | -1.20 |
| Species | Cellulosilyticum uncultured bacterium | 8.10e-3 | 0.67 |

Table S6. Results of the Kruskal Wallis test comparing relative abundance of taxonomic classifications between June-July, June-August, and July-August for Caw Ridge: a total of 0/13 phyla, 3/18 classes, 0/22 orders, 4/36 families, 3/78 genera, and 3/82 species classifications. The difference in percent relative abundance is shown (relative abundance of June–relative abundance of July; relative abundance of June–relative abundance of August; or relative abundance of July–relative abundance of August).

| Level | Comparison | Classification | p-value | % Diff Rel Abun |
| --- | --- | --- | --- | --- |
| Class | July-August | Coriobacteriia | 1.96e-3 | -2.44e-2 |
| Class | July-August | Clostridia | 1.14e-3 | -1.32e-1 |
| Class | July-August | Negativicutes | 3.43e-3 | 4.03e-3 |
| Family | July-August | Christensenellaceae | 2.98e-3 | -7.20e-2 |
| Family | July-August | Clostridiales Family XIII | 1.22e-3 | -6.21e-2 |
| Family | June-August | Lachnospiraceae | 8.89e-4 | -8.85e-3 |
| Family | June-August | Ruminococcaceae | 1.86e-3 | 3.09e-2 |
| Genus | July-August | Clostridiales Family XIIIAD3011 group | 186e-3 | -2.41e-2 |
| Genus | July-August | Erysipelatoclostridium | 2.43e-3 | 8.34e-3 |
| Genus | July-August | Eubacterium hallii group | 5.24e-3 | -4.03e-2 |
| Genus | June-July | Eubacterium hallii group | 1.42e-3 | 4.11e-2 |
| Species | July-August | Erysipelatoclostridium uncultured bacterium | 1.86e-3 | 1.45e-2 |
| Species | July-August | Eubacterium hallii group uncultured bacterium | 1.59e-3 | -3.51e-2 |
| Species | June-August | Erysipelatoclostridium uncultured bacterium | 8.48e-3 | 1.34e-2 |
| Species | June-July | Eubacterium hallii group uncultured bacterium | 1.59e-3 | 3.51e-2 |
| Species | July-August | Ruminococcaceae NK4A214 group uncultured bacterium | 1.53e-3 | 4.31e-2 |
